# Anoxia Tolerant DNA Replication is Supported by ATR Kinase in the Annual Killifish *Austrofundulus limnaeus*

**DOI:** 10.64898/2026.06.01.729397

**Authors:** Riley Roth-Carter, Erin Helms, Joshua C Saldivar, Jason Podrabsky

## Abstract

Hypoxia and anoxia are known to suppress cell proliferation due to an increase in replication stress and activation of DNA damage checkpoints. Embryos of the annual killifish *Austrofundulus limnaeus* show a strong tolerance to extended anoxic exposure, indicating an improved genomic stability under oxygen starvation. Here we investigate the cell cycle regulation of the anoxia tolerant killifish embryonic cell line PSU-AL-WS40NE during anoxic exposure. Live cell imaging confirms continued cell proliferation of WS40NE cells for the first 24 hours of anoxic exposure with minimal cell death. Fluorescent imaging shows that cells begin to accumulate in G_1_ after the first day in anoxia with a pronounced and rapid entry into the S phase upon reoxygenation. Pharmacological inhibition tests show that this response appears to be reliant more on ATR signaling then ATM, suggesting that increased γH2AX levels are driven by increased replication stress instead of DNA damage. This conclusion is further supported by an apparent lack of induction of a G_2_ checkpoint in these cells suggesting that DNA damage during anoxic replication is minimal. Maintaining cellular proliferation during initial exposure to anoxia and accumulating cells in the G_1_ phase for extended anoxic exposure is likely one way that embryos of the annual killifish are able to survive prolonged anoxia and provides insight into mechanisms that enable cells to proliferate under metabolic stress.

## Introduction

Maintaining a stable genome is a vital requirement for cell and organismal survival and proliferation, with errors in processes involved in genome maintenance potentially leading to disease, dysfunction, and death (Tomasetti et al., 2017; Zeman and Cimprich, 2014). Because of the potentially disastrous consequences of passing on genomic damage to daughter cells, there are numerous DNA damage and cell cycle checkpoints that function to facilitate genome repair prior to progressing through the cell cycle (Boddy et al., 1998; Elledge, 1996). Cells are routinely able to withstand these basal levels of genotoxic stress but when the cellular environment changes, such as within hypoxic tumor microenvironments, genome maintenance pathways can become strained and unable to resolve high levels of damage that can accumulate in these environments (Riffle et al., 2017). Detection of DNA damage by ataxia telangiectasia mutated (ATM) and ataxia telangiectasia and Rad3 related (ATR) proteins can lead to cells stopping at the G_1_/S checkpoint (Abraham, 2001). ATM protein is activated in response to double stand breaks in the DNA while ATR is activated in response to replication stress (Awasthi et al., 2015). Both proteins activate checkpoint kinases 1 and 2 (CHK1/2) and stabilize levels of p53 to induce cell cycle arrest through the up regulation of p21 (El-Deiry et al., 1993). With DNA damage directly leading to exit from the cell cycle, it is expected that exposure to genotoxic stressors will cause a halt in the cell cycle until the stressor is removed and damage repaired. This strong control of the cell cycle prevents cell proliferation when DNA damage is present, and it also confirms replication does not continue when cells are exposed to stressful environments. Tolerance to high levels of environmental genotoxic stressors is a hallmark of metabolic dormancy in animals, as well as for radio-resistant human cancers and thus is an important area for continued research.

The effect of hypoxia and anoxia on control of the cell cycle is not completely understood. Hypoxia has been shown to activate p53 protein and leads to increased expression of p21 and p27 proteins, core proteins that inhibit cell proliferation, leading to cell cycle arrest (Druker et al., 2021; Hubbi and Semenza, 2015; Koshiji et al., 2004). At the same time, hypoxia has also been shown to cause an upregulation in aurora A (AURKA), a mitotic kinase, causing hypoxia-induced cellular proliferation (Klein et al., 2008). Cell proliferation during exposure to stressful conditions, especially hypoxia and anoxia, is known to lead to high mutational burden and in cancers is a driver of malignant progression (Vaupel, 2004).

Embryos of the annual killifish are extremely tolerant to many different known genotoxic stressors, such as high dose UV-C and ionizing radiation, 3% H_2_O_2_, and months of complete anoxia (Podrabsky et al., 2016; Podrabsky et al., 2007; Wagner et al., 2019; Wagner and Podrabsky, 2015). These conditions are known to cause DNA damage and should induce withdrawal from the cell cycle based on data from mammalian systems. Primary cells that have been isolated from embryos of the annual killifish in normal conditions appear to be mostly in the G_1_ phase of the cell cycle and presumably quickly pass through S and G_2_. This is in stark contrast to embryos from other fish species where many cells are found in S and G_2_ (Meller et al., 2012). When the embryos are exposed to 24 hours of anoxia there is no effect on the percentage of cells within G_1_ and G_2_ when compared to normoxic conditions (Meller et al., 2012). This accumulation of cells in G_1_ is hypothesized to allow for decreased energy cost for genome maintenance during prolonged anoxic exposure. This signature could be caused by cells immediately arresting at the stage they are in when exposed to anoxia, as seen in other stress tolerant organisms such as the embryos of the soil nematode *Caenorhabditis elegans* that also enter into a dormant state and exit from the cell cycle when exposed to anoxic conditions (Hajeri et al., 2005). Embryos of the zebrafish, *Danio rerio*, can also arrest when exposed to anoxia, but seem to only arrest in the S and G_2_ phases of the cycle (Padilla and Roth, 2001). The high proportion of G_1_ phase cells in *A. limnaeus* embryos suggests a potentially unique regulation of the cell cycle during development and in response to stress. However, previous studies relied on DNA content alone to define cell cycle kinetics, limiting the ability to draw more mechanistic conclusions concerning cell cycle regulation in response to anoxia.

To better study cellular mechanisms associated with this anoxia tolerance, in this study we use an immortal cell line isolated from late-stage *A. limnaeus* embryos, PSU-AL-WS40NE (Riggs et al., 2019). This cell line exhibits a strong tolerance of anoxia, with many cells surviving 7 weeks without oxygen at 30°C (Riggs et al., 2019). Along with this impressive anoxia tolerance these cells appear to continue to proliferate for several days when exposed to anoxia based on cell counts (Riggs et al., 2019). Continuing to replicate DNA and proliferate while exposed to these genotoxic stressors would presumably cause large levels of DNA damage and cell death, as is seen in other systems (Vaupel, 2004). Yet, WS40NE cells exposed to anoxia and reoxygenation can resume normal cellular functions and proliferation. Discerning the ability of the annual killifish to continue cell proliferation during anoxic exposures without deleterious effects, shown by continued normal embryonic development after extended anoxia and reoxygenation, can lead to a better understanding of the role of cell cycle regulation in the extreme anoxia tolerance of these cells and may help determine the drivers of radio- and chemo-resistance in cancers that exhibit similar cell stress responses to oxygen deprivation (Podrabsky et al., 2007).

## Results

### PSU-AL-WS40NE continue Cellular Replication during Anoxia and Reoxygenation

To confirm and expand on previous data suggesting that the cell line PSU-AL-WS40NE (WS40NE) is able to continue proliferation during initial anoxic exposure, live-cell bright field images were collected over a time-course of 24 hours of anoxia followed by 24 hours of reoxygenation to visualize cellular morphology, behavior, and divisions. These images show clear cell divisions occurring during anoxia and survival of the daughter cells throughout the anoxic exposure (Fig. 1a). Daughter cells produced during anoxia are also able to divide again following reoxygenation (Fig. 1a). Cell counts of these images reveal that cellular proliferation continues throughout the entire 24 hours of anoxia at similar rates to cells in normoxia, suggesting that there is no pause in the cell cycle within the first 24 hours of anoxia. This trend continues through reoxygenation as well, though there may be a slight depression in early time points of reoxygenation followed by a slight increase in overall proliferation after 24 hours of reoxygenation compared to 48 hours of normoxia (Fig 1b). Based on cellular morphology, less than 2% of cells experience cell death during the time-course of anoxia and aerobic recovery (Fig. 1c).

**Fig 1.**
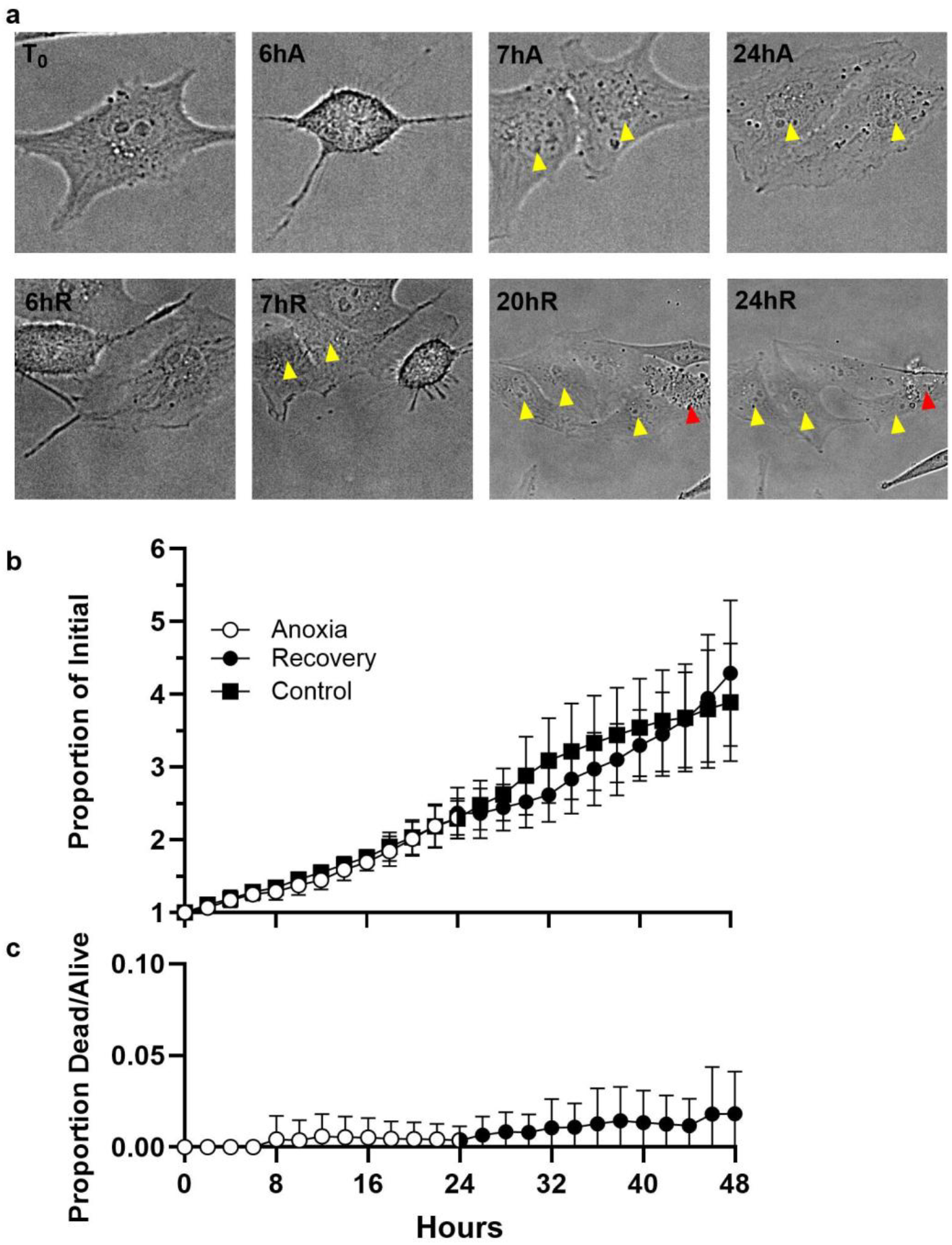
Live Bright Field Imaging of WS40NE Cell during Anoxia and Reoxygenation. (a) A single cell followed through both anoxia and reoxygenation. Yellow arrows indicate surviving daughter cells and red arrows indicate morphologically dead daughter cells. (b) Quantification of cell counts during anoxia and reoxygenation and a normoxic control as a proportion of the initial cells. Results presented as mean ± SD, n = 3. (c) Quantification of the proportion of morphologically dead cells compared to live cells at that same timepoint. Results presented as mean ± SD, n = 3.

### Anoxia Induces Delayed G_1_ Arrest with Rapid and Sustained S-Phase Increase after Reoxygenation

With evidence of continued cell proliferation during short exposures to anoxic conditions, we wanted to determine the effect of longer anoxic exposure and get improved resolution into the cell cycle during anoxia. To do this we utilized Quantitative Image Based Cytometry (QIBC) with EdU pulsing to quantify cell cycle dynamics across a time course of anoxia and reoxygenation in WS40NE cells (Luis et al., 2013). We also monitored levels of γH2AX, a marker for DNA damage and replication stress. EdU incorporation is significantly reduced after 24 hours of anoxia and by 96 hours almost all EdU incorporation has stopped (Fig 2). Interestingly we see that after only 4 hours of reoxygenation there is already a population of cells restarting S-phase, as indicated by EdU incorporation, and by 24 hours we see a return to normal levels of EdU incorporation in S phase cells. Analysis of cell cycle phases revealed normoxic cells to have relatively equal levels of G_1_ and S-phase cells, with 49.6% and 45.7% respectively, and only a small proportion (4.7%) of cells in G_2_. As cells grow towards confluence in normoxia, there is a slow trend towards an increase in G_1_ phase and decrease in S-phase cells until there is a majority, 60.4%, of cells in the G_1_ phase of the cell cycle with 28.6% and 11% of cells in S and G_2_, respectively, after 32 h (Fig. 3a). This end timepoint of normoxia is in line with previous analysis of whole embryos of the annual killifish which found around 80% of cells in G_1_ (Meller et al., 2012). Interestingly, after initial exposure to anoxia there is a stabilization or perhaps even a slight increase in percentage of S phase cells after 4 h, suggesting sustained and perhaps even increased DNA replication during the initial transition into anoxia (Fig 3b). The number of cells actively replicating their DNA (in S phase) then slowly declines reaching a low around 20% after 48 h and 12% after 96h of anoxia (ANOVA, p = 0.0009 and < 0.0001 respectively). As S phase cells decrease, there is an increase in G_1_-phase cells to around 70% of cells after the first 48 h and 77% by 96 h of anoxia (ANOVA, p = 0.0025 and 0.0004 respectively). There is little change in the level of cells in G_2_ at 96 h of anoxia compared to normoxic levels, comprising the remaining 12% of cells (ANOVA, p = 0.001).

**Fig. 2.**
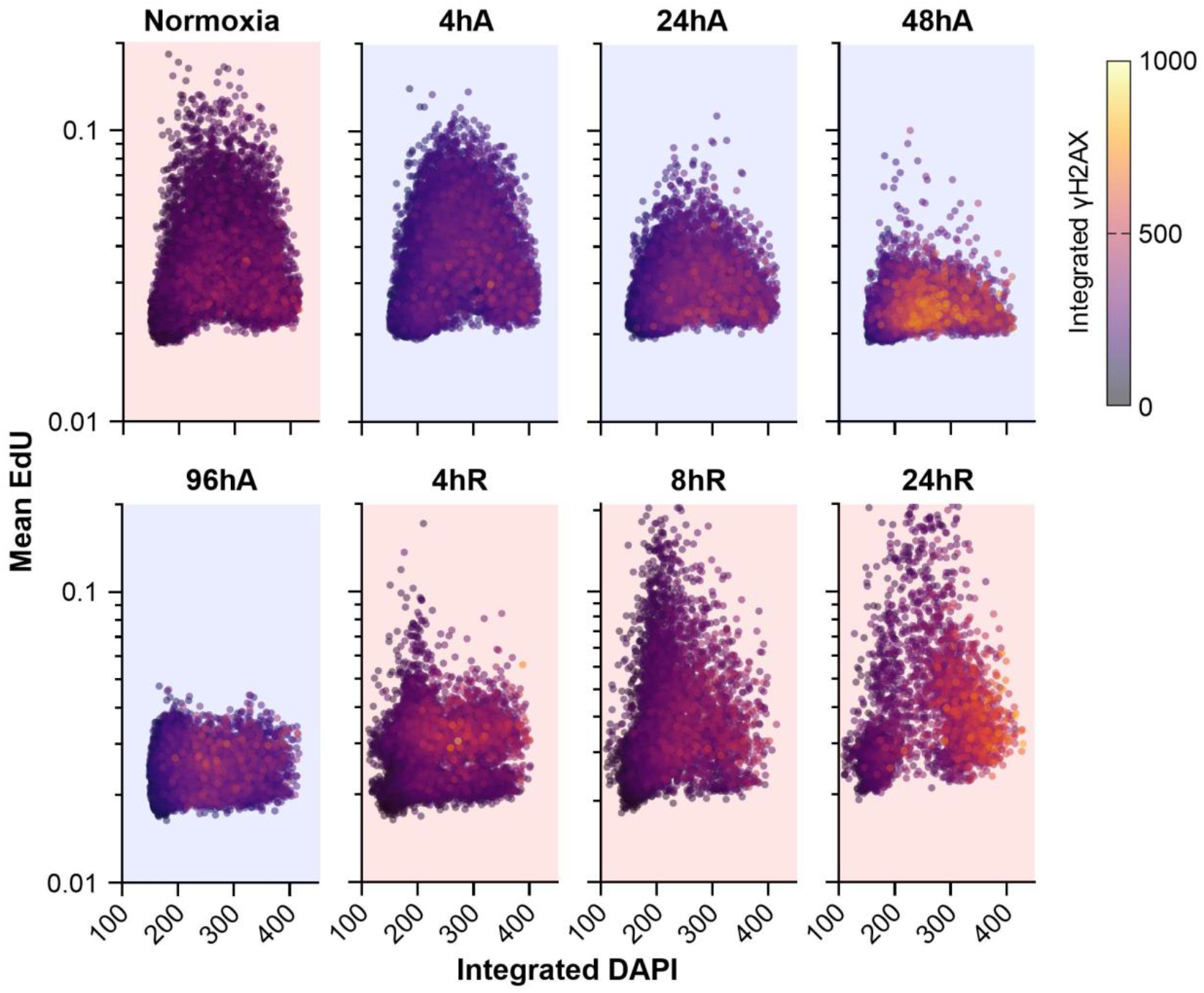
Quantitative Image-Based Cytometry data of WS40NE Cells during Anoxia and Reoxygenation. WS40NE cells were plated at 1.5 × 10^5^ cells per well in a glass-bottom optical 96 well plate and then exposed to anoxic conditions for 96 hours followed by 24 hours of normoxic conditions. Each dot represents an individual cell’s EdU and DAPI fluorescence with dot color indicating levels of γH2AX. Red background indicates normoxic control and reoxygenation, blue background indicates anoxic conditions.

**Figure 3.**
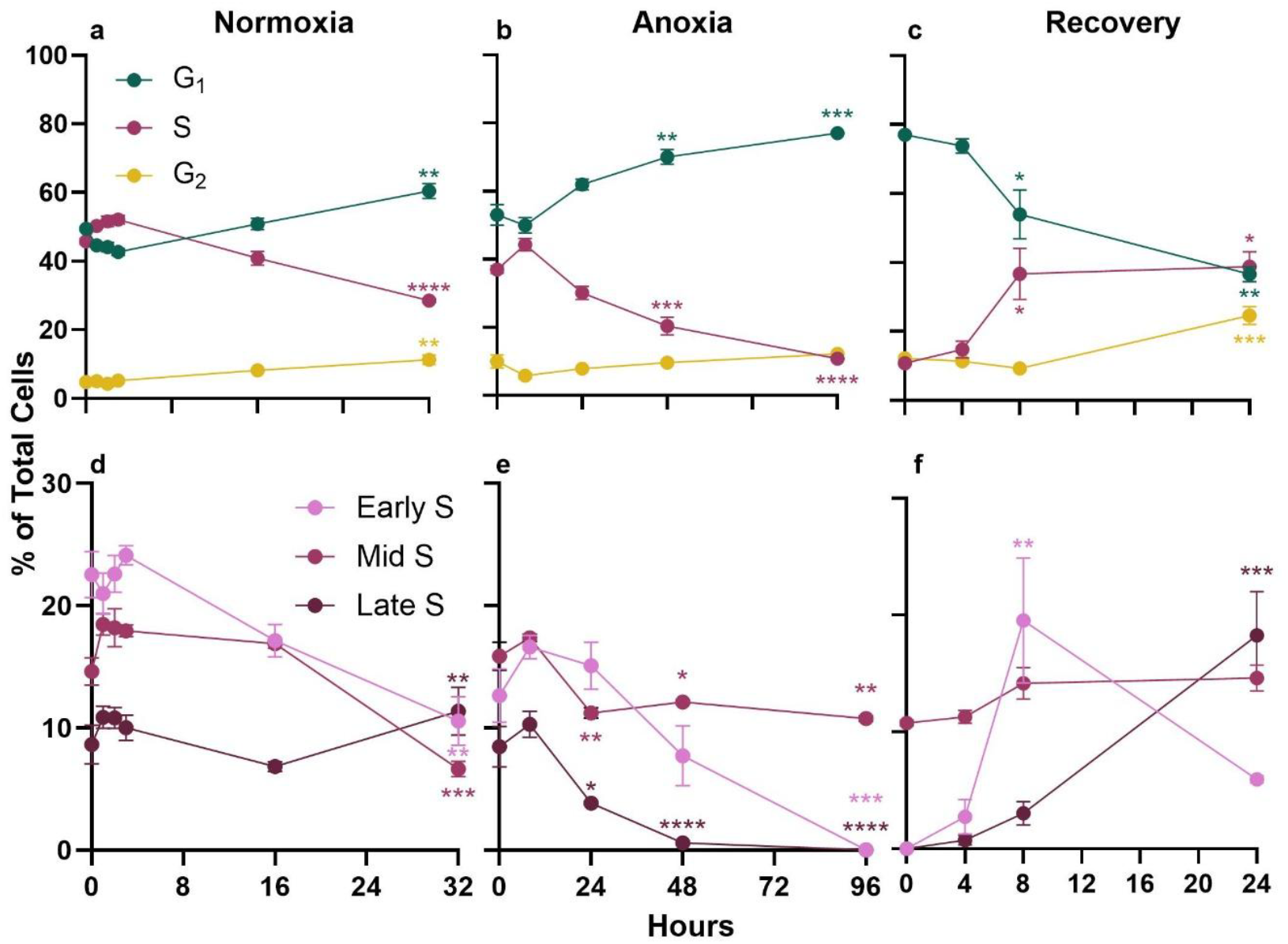
Cell Cycle Analysis of WS40NE Cells Exposed to Anoxia and Reoxygenation. Proportion of cells within each stage of the cell cycle were quantified during anoxia and reoxygenation. Proportion of each stage of the cell cycle in cells exposed to: (a) normoxia for 32 hours or (b) anoxia for 96 hours followed by (c) reoxygenation for 24 hours. (d-f) The proportion of cells in early-. Mid-, and late-S phases during (d) normoxia, (e) anoxia and (f) reoxygenation. Results presented as mean ± SEM, n = 3. Significance was measured using One-Way ANOVA, with Dunnett’s post hoc test comparing all time points within a single stage to the 0 hour point within that condition, p < 0.05 *, p < 0.01 **, p < 0.001 ***, p < .0001 ****.

After 4 days of anoxia we reintroduced cells to normoxic conditions to observe the effect of reoxygenation on cell cycle dynamics. These data, coupled with the low number of dead cells reported in Figure 1, support a quick re-entry of cells into the cell cycle and resumption of DNA replication after 8 h of aerobic recovery (Fig. 3c; ANOVA, p = 0.003). This increase comes with a comparable decrease in the G_1_ population (p = 0.03) at 8 h. This large increase in S-phase cells and comparable decrease in G_1_ is maintained for the first 24 h of recovery. Interestingly, there is also a significant increase in G_2_ cells after 24 hours of recovery (p = .003). If S phase cells are divided into early, mid, and late based on DNA content (Fig. 3d,e,f), a marked increase in early S-phase cells is observed after 8 h of aerobic recovery (Fig. 3f; p = 0.003). These data are consistent with cells entering S phase after arrest at the G_1_/S checkpoint during anoxic exposure.

### Extended Anoxic Exposure causes Increased γH2AX

This continuation of proliferative cell cycle dynamics for the first 48 h of anoxia is intriguing in the context of increased levels of γH2AX, a marker of both replication stress and DNA damage (Figs. 2,4a). At 48 hours of anoxia there is a significant increase in γH2AX in all stages of the cell cycle. Interestingly by 96 hours of anoxia this increase in γH2AX completely lost. During recovery the only significant increase in γH2AX is seen at 24 hours of recovery and only in Late-S and G_2_ cells.

**Figure 4.**
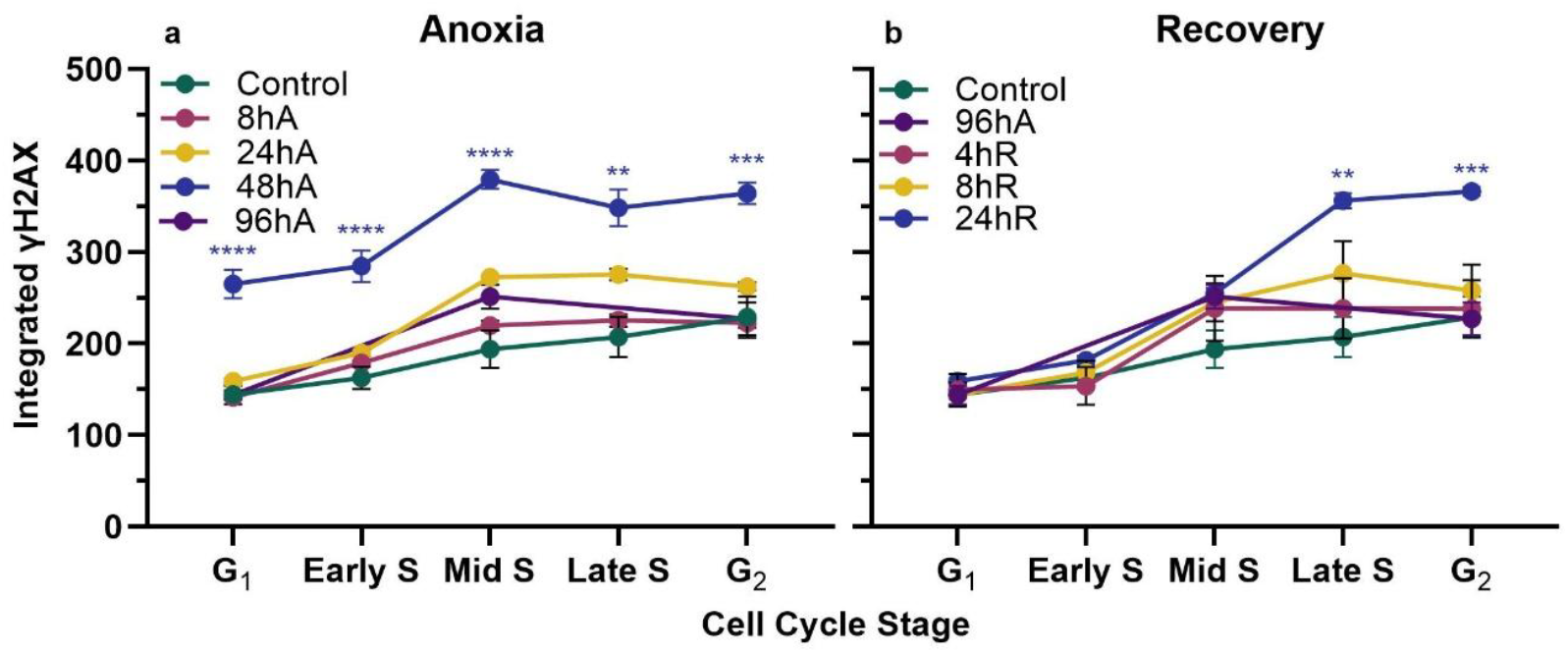
Quantification of γH2AX during anoxia and reoxygenation across Cell Cycle Stages. Integrated γH2AX signal from QIBC images taken across (a) 48 hours of anoxia and (b) 24 hours of reoxygenation. Results presented as mean ± SEM (n = 3). Significance was measured using One-Way ANOVA and Dunnett’s post hoc test comparing all time points within a single cell cycle stage to control within that condition, p < 0.05 *, p < 0.01 **, p < 0.001 ***, p <0.0001 ***.

### Increased γH2AX is Driven by Replication Stress

As the increase of γH2AX seen during anoxia can either be attributed to an increase in replication stress or DNA damage, we used small molecule inhibitors of the key kinases ATR and ATM to test for the source of γH2AX. During anoxia and normoxia ATM has no clear effect on either the distribution of cells across the cell cycle or the intensity of γH2AX (Fig. 5 and 6). By contrast, the inhibition of ATR causes an increase in G_1_ phase cells during normoxia, and an increase in γH2AX intensity in G_1_ (Fig. 5). These increases are likely due to the mitotic transmission of under-replicated DNA to the daughter G_1_ cells (Chanoux et al., 2009; Guo et al., 2000; Pedersen et al., 2015). Strikingly, ATR inhibition under anoxia causes a dramatic increase in γH2AX in S phase cells (Fig. 6), suggesting anoxia tolerance is dependent on ATR signaling to prevent replication fork collapse and nucleus-wide DNA damage. Taken together this suggests that anoxia does not cause significant amounts of DNA damage in S phase, but rather increases replication stress which is addressed by ATR signaling to prevent global replication fork collapse.

**Figure 5.**
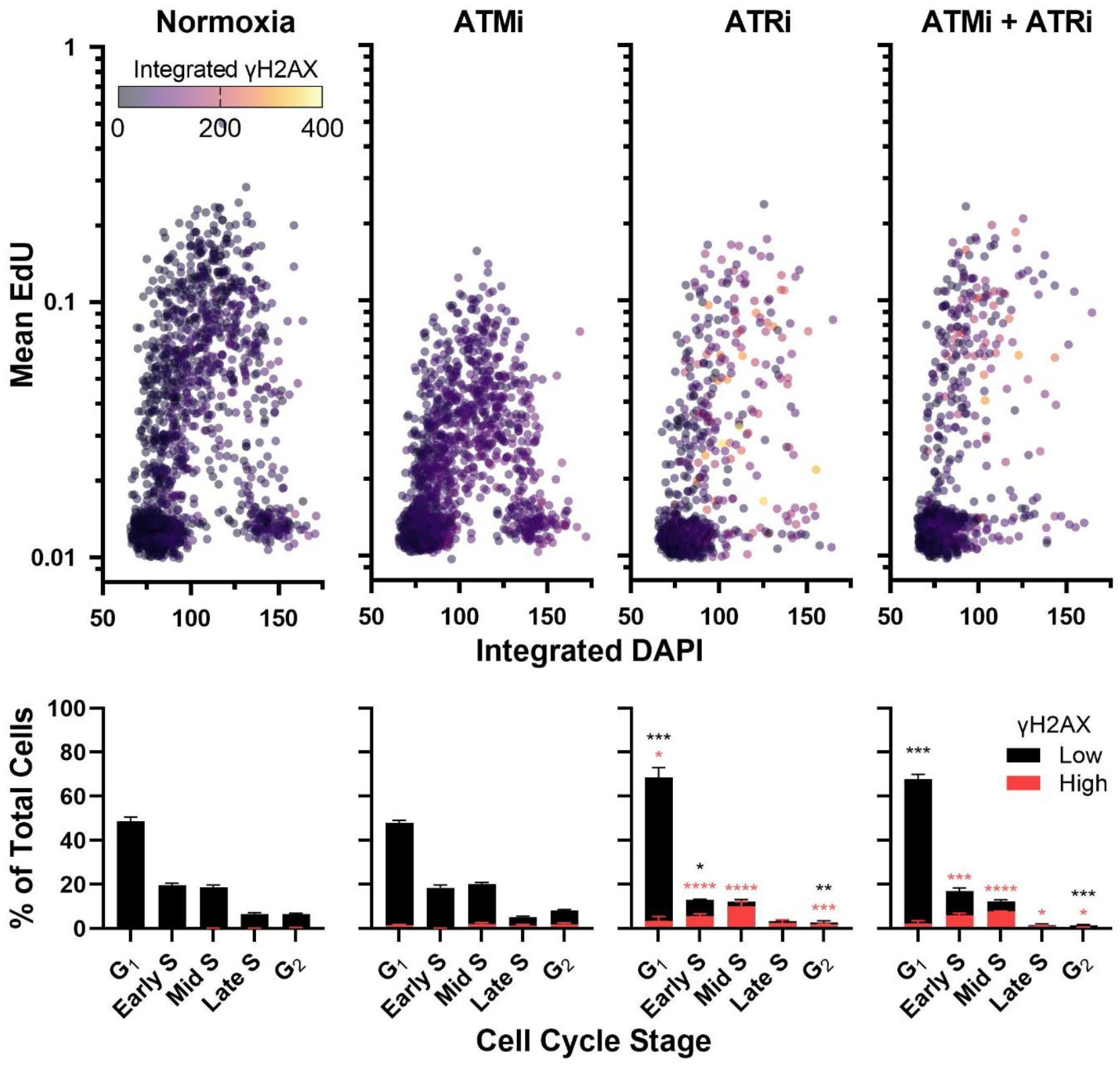
Effect of Phosphatidylinositol 3-kinase-related kinases (PIKKs) inhibitors on the cell cycle and γH2AX. WS40NE cells were exposed to inhibitors of ATR (8 µM AZ20) and ATM (0.01 µM AZD0156) in normoxic conditions for 24 hours. (Top panels) Each dot represents an individual cell’s EdU and DAPI fluorescence with dot color indicating levels of γH2AX. (Bottom panels) Each cell cycle stage quantified as proportion of total cells and the proportion of high γH2AX cells within each stage. Cells are indicated as γH2AX high if integrated fluorescence intensity is higher than normoxic controls. Results presented as mean ± SD (n = 3). Significance was measured using One-Way ANOVA and Dunnett’s post hoc test comparing proportions of each cell stage and high vs. low γH2AX to control, p < 0.05 *, p < 0.01 **, p < 0.001 ***, p < 0.0001 ****. Black asterisks represent changes in cell cycle stage proportion and red asterisks represent changes in proportion of high vs. low γH2AX.

**Figure 6.**
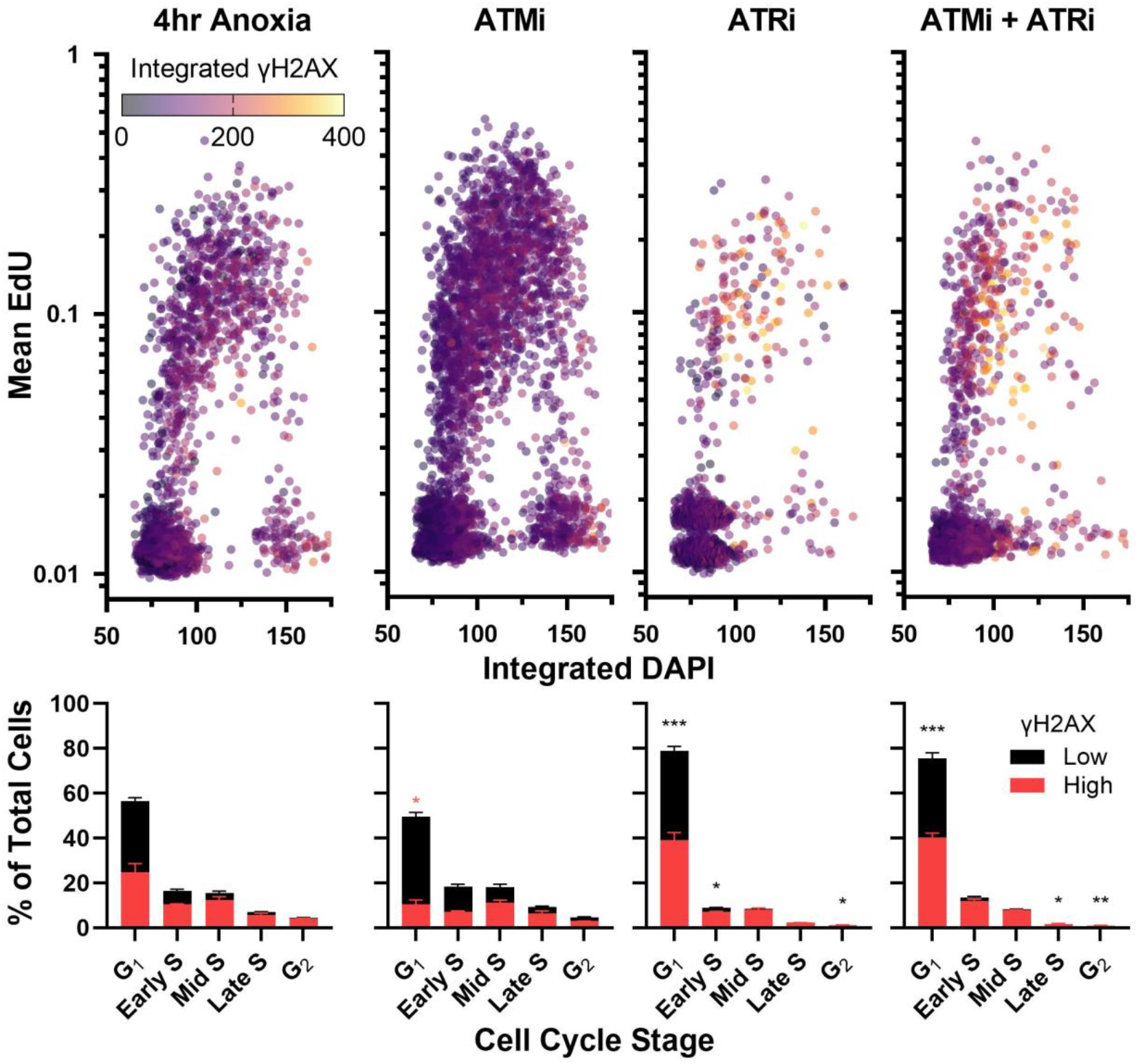
Inhibition of ATR affects cell cycle kinetics during anoxia. WS40NE cells were exposed to inhibitors of ATR (8 µM AZ20) and ATM (0.01 µM AZD0156) for one hour of normoxia and then 4 hours of anoxia. (Top panels) Each dot represents an individual cell’s EdU and DAPI fluorescence with dot color indicating levels of γH2AX. (Bottom panels) Each cell cycle stage quantified as proportion of total cells and the proportion of high γH2AX cells within each stage. Cells are indicated as γH2AX high if integrated fluorescence intensity is higher than normoxic control. Results presented as mean ± SD (n = 3). Significance was measured using One-Way ANOVA and Dunnett’s post hoc test comparing proportions of each cell stage and high vs. low γH2AX to control, p < 0.05 *, p < 0.01 **, p < 0.001 ***, p < 0.0001 ****. Black asterisks represent changes in cell cycle stage proportion and red asterisks represent changes in proportion of high vs. low γH2AX.

## Discussion

Here we report that the WS40NE cell line transitions into anoxia without major disruptions to cell cycle progression or DNA replication. Rather, they react through a slow and controlled process that results in cells accumulating in G_1_. This is a unique pattern when compared to other vertebrate systems. Hypoxia and anoxia typically induce a rapid arrest in cell proliferation, likely an adaptive response to prevent increased cell populations and thus increased oxygen demand within tissues, with continued hypoxic exposure leading to cell death (Baldea et al., 2018; Carmeliet et al., 1998; Gardner et al., 2001; Goda et al., 2003; Hubbi and Semenza, 2015). However, this general response is not universal with many cancers and stem cell populations showing increased cell proliferation during hypoxic exposure (Gordan et al., 2007; Hubbi and Semenza, 2015; Morrison et al., 2000).

Continued cellular proliferation with slow loss in S-phase cells during anoxic exposure in WS40NE cells is similar to what is observed in the gills of the anoxia tolerant Crucian carp, *Carassius carassius* (Sollid et al., 2005). Though in the Crucian carp there is not a rapid increase in S phase cells after reoxygenation. However, it is worth noting that at the organismal level killifish embryos and Crucian carp have differing strategies in response to anoxia. Carp remain active during anoxia and rely heavily on temperature-driven metabolic depression and glycolytic energy production, while killifish embryos enter a profound state of metabolic and developmental dormancy. Thus, the continued cell proliferation in killifish cells may be viewed as an adaptive mechanism to support reversible cell cycle arrest, rather than a continued proliferation supported by changes in metabolic pathways.

In most vertebrate cells, mismatch between ATP production and consumption during anoxia leads to an energetic crisis and induction of cell death (Taylor and Pouyssegur, 2007). In contrast, during reoxygenation an increase in cellular reactive oxygen species (ROS) is the driver of cell death (Eleftheriadis et al., 2020; Neri et al., 2017). In WS40NE cells, exposure to anoxia and reoxygenation do not cause high levels of cell death, especially when compared to other vertebrate models (Azad et al., 2008; Carmeliet et al., 1998; Lefevre et al., 2017; Zhu et al., 2010). Thus, WS40NE cells show a remarkable capability to survive without oxygen and to maintain viability through reoxygenation. There is evidence that cancer cells respond to hypoxic exposure by preparing for reoxygenation, which then assists in rapid recovery and cellular proliferation after they return to normoxic conditions (Schlaepfer et al., 2015). It is likely that the WS40NE cells are responding in a similar way, evidenced by the rapid increase of S phase cells beyond that observed in normoxia as an early response to reoxygenation. In whole embryos this rapid proliferation after reoxygenation could be vital to repair damaged tissues or could be a compensatory response to increase developmental rate post-anoxia.

The ability to maintain cellular proliferation during hypoxic and anoxic exposures requires functioning DNA damage repair pathways. *H2afx* ^*-/-*^ mice, a deletion of the gene encoding for histone H2AX and thus required for the production of γH2AX, lose hypoxia-driven endothelial cell proliferation (Economopoulou et al., 2009). In most cells, hypoxia is a known driver of replication stress and the DNA repair response is required to efficiently restart stalled replication forks (Ng et al., 2018). Consistent with this, our data show that the replication stress response kinase, ATR, protects S phase cells from accumulating DNA damage allowing DNA replication for several hours in anoxic conditions. However, WS40NE cells eventually reduce replication. Extended anoxia causes a G_1_ arrest and a return of γH2AX levels to near normoxic levels after 4 d of anoxia. This may suggest the completed repair of DNA damage and the accumulation of cells that are prepared to proliferate almost immediately upon reoxygenation. Notably, even with both ATR and ATM inhibition during anoxia exposure, high levels of γH2AX are observed. This suggests activation of the third protein in the family of kinases that can phosphorylate H2AX, DNA-PKcs (An et al., 2010).

In conclusion, WS40NE cells are able to maintain normal cellular proliferation for up to 48 hours of anoxic exposure, even with apparent increased replication stress. This anoxia tolerance is associated with sustained DNA replication for several hours and a reliance on the replication stress response kinase ATR to prevent widespread DNA damage in S phase. Continued proliferation potentially allows the cells to reach a stage of the cell cycle that promotes survival of extended anoxic exposures and prepares them for recovery during reoxygenation. Anoxia and reoxygenation are an integral part of several different pathological states and environmental conditions, and better understanding the molecular mechanisms that support WS40NE cells to survive these stressors holds promise for the development of new interventions and therapies.

## Methods

### PSU-AL-WS40NE Cell Culture

The PSU-AL-WS40NE cell line is a neuroepithelial line derived from Wourms’ Stage 40 embryos of *Austrofundulus limnaeus* (Riggs et al., 2019). Cells were maintained in complete cell culture media comprised of Gibco ^™^ Leibovits’s L-15+Glutamine (11-415-114, ThermoFisher Scientific, Waltham, MA, USA), 8.5% fetal bovine serum (FBS; 97068-085, VWR, Radnor, PA, USA), 5mM glucose (45001-116, VWR, Radnor, PA, USA) and 100U/mL penicillin/streptomycin (pen/strep; 15-140-122, ThermoFisher Scientific, Waltham, MA, USA) in 100 × 20 mm CytoOne^™^ Tissue Culture dishes (CC7682-3394, USA Scientific, Ocala, FL, USA) at 30°C under normal atmospheric conditions. For experiments, cells were grown in 96 well optical culture plates designed to be used for imaging (P96-1.5P, Cellvis, Mountain View, CA, USA).

### Anoxic Exposure

Anoxic cell culture media was prepared as previously described (Riggs et al., 2019). Briefly, pen/strep and glucose were added to L-15 to a final concentration of 100U/mL and 5mM, respectively. The required amount of FBS to reach a final concentration of 8.5% was injected into a sterile hydrated Slide-A-Lyzer^™^ dialysis cassette (A52964, ThermoFisher Scientific, Waltham, MA, USA) using a sterile syringe and 22 gauge needle. The cassette was then placed within the L-15 media and bubbled for 1 hr with sterile-filtered (0.22 μm) high purity nitrogen gas before being transferred into an anaerobic chamber (Bactron EZ, Sheldon Laboratories, Cornelius, OR, USA) with an atmosphere of 5% H_2_ and 95% N_2_. The media was bubbled within the chamber overnight with 0.22 μm filtered chamber gas mixture using an aquarium pump. The following day, the FBS was withdrawn from the cassette with a sterile needle and syringe and added to the L-15 media.

The complete media was again sterile filtered (0.22 μm) within the chamber. Anoxic phosphate buffered saline (PBS) was generated by bubbling with nitrogen gas as above followed by overnight equilibration to the atmosphere in the anaerobic chamber.

To expose cells to anoxia, normoxic cell culture media was removed in a biosafety cabinet, the 96 well plate was transferred into the anaerobic chamber through a pass box, and then the cells were washed three times with anoxic PBS. After the washes, cells were bathed in 200 μl per well of anoxic cell culture media. Cells were maintained at 30°C using the anaerobic chamber’s built-in incubator. For recovery samples, cells were removed from the anaerobic chamber and washed three times with normoxic 1X PBS followed by incubation in normoxic complete cell media in an incubator at 30°C under normal atmospheric conditions.

For some experiments it was essential to have normoxic and anoxic samples in the same 96-well plate for imaging. In these circumstances, normoxic conditions were maintained in wells while the plates were within the anoxic chamber by sealing them with aluminum foil plate seals (FCS-25, AlumaSeal® CS^™^, Excel Scientific, Victorville, CA, USA) under normoxic conditions with 100 μL of normoxic complete cell media just prior to placing the plate into the anaerobic chamber. Sealed normoxic wells maintained aerobic conditions (80% saturation with O_2_ or greater) for up to 24 h when sealed in this manner and placed within the anaerobic chamber (data not shown).

### Immunofluorescence staining

Immunofluorescence staining and imaging were performed as recently described (Marmolejo et al., 2026). Briefly, cells were grown on 96-well optical plates (Cellvis, P96-1.5P) with a seeding density of 1.5 × 10^4^ cells/well. Cells were exposed to experimental conditions, including anoxia, 2 mM hydroxyurea (ab142613, Abcam, Waltham, MA, USA), 8 µM AZ20 (an inhibitor of ATR; HY-15557, MedChemExpress, Monmouth Junction, NJ, USA), or 0.01 µM AZD0156 (an inhibitor of ATM; HY-100016, MedChemExpress, Monmouth Junction, NJ, USA). If cells were exposed to inhibitors, they were given a pre-dose of inhibitors for 1h prior to being moved to anoxic conditions and continually exposed to the drug during anoxia. During the final 15 min of exposure, cells were exposed to a pulse of 20 µM 5-ethynyl-2’-deoxyuridine (EdU; 900584, Sigma-Aldrich, St. Louis, MO, USA). After exposure to EdU, cells were washed 3 times with 1X PBS and fixed with 4% paraformaldehyde (PFA) in PBS for 15 min. Fixed cells were stored at 4°C in PBS for up to 24 h before staining. Cells were permeabilized using ice-cold methanol for 10 min at 4°C and washed 3 times with 1X PBS at room temperature. Cells were incubated in 3% bovine serum albumin (BSA; A3294, Sigma-Aldrich, St. Louis, MO, USA) in 1X PBS for 10 min prior to initiation of the Click-iT reaction for EdU fluorophore conjugation using the Click-iT Cell Reaction Buffer kit (C10269, Invitrogen, Carlsbad, CA, USA) and Alexa Fluor™ 647 Azide (1 µg/ml; A10277, Invitrogen, Carlsbad, CA, USA) according to the manufacturer’s guidelines. Cells were incubated for 30 min at room temperature with constant rocking. After the Click-iT reaction, cells were washed with 3% BSA/PBS, followed by 3 washes in 1X PBS. Cells were blocked with 1% BSA in PBS for 1 h at room temperature on a bench top rocker. Following blocking, cells were incubated in rabbit polyclonal anti-zebrafish H2A.X phospho-Ser139 (1:500; 89424-450, VWR, Radnor, PA, USA) overnight at 4°C with constant agitation. Following the overnight incubation with primary antibodies, cells were washed 3 times in 1X PBS at room temperature and co-stained with 5 μg/mL DAPI (ab285390, Abcam, Waltham, MA, USA) and an anti-rabbit Alexa Fluor 488 conjugated antibody in 1% BSA (1:1000; ab150077, Abcam, Waltham, MA, USA) for 1 h at room temperature on a bench rocker. Finally, cells were washed three times with 1X PBS and imaged in 1X PBS.

### Quantitative image-based cytometry and analysis

Immuno-stained cells were imaged on a fully-automated ImageXpress Micro Confocal Imaging System (Molecular Devices, San Jose, CA, USA) following methods previously described (Marmolejo et al., 2026). Widefield images were captured using a 20X Ph1S Plan Fluor ELWD ADM 0.45 NA objective and an sCMOS camera. Following image acquisition, CellProfiler (Broad Institute, Cambridge, MA, USA) was used for Image analysis (Stirling et al., 2021). DAPI-stained images were used to create a nuclear mask for nuclear intensity measurements, and the mean intensity values for each marker were measured within the nuclear mask. DNA content was determined using DAPI integrated intensity.

### Live Cell Imaging

Cells were grown on 96-well optical plates (Cellvis, P96-1.5P) with a seeding density of 1.5 × 10^4^ cells/well. Cells were then transferred to anoxic conditions as described above in 100µL of anoxia media. To maintain anoxic conditions during live imaging two drops of anoxic mineral oil were floated on top of each well using a sterile disposable pipette before sealing with transparent sealing tape. This maintained anoxic conditions for at least 24 h while the plate was outside of the anaerobic chamber (data not shown). Brightfield images were taken every 15 min for 24 h on an Olympus IX83 inverted research microscope (Evident Scientific, State College, PA, USA). Plates were then removed from the microscope and returned to normoxic conditions by pipetting normoxic media into the bottom of the well and washing out the mineral oil by overflowing the wells with normoxic media. This floated the anoxic mineral oil up and out of the wells and prevented exposing cells to oil which made imaging difficult. The plates were placed back onto the microscope and recovery from anoxia was documented with images taken every 15 min for 24 h. Final images were processed in ImageJ where the number of total cells and morphological dead cells were counted at 2-hour timepoints. (ImageJ 1.54p, National Institutes of Health, USA). Dividing cells and their daughter cells were tracked through exposure to anoxia and through reoxygenation and recovery.

### Statistical Analysis

For all comparisons of proportional data, values were normalized and transformed through the arcsine square root transformation 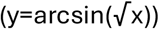 before One-Way ANOVA with Dunnett’s post hoc test used to determine significance between means of individual treatments compared to normoxic controls.

